# Development of an HPTLC method for determination of hypoglycin A in aqueous extracts of seedlings and samaras of *Acer* species

**DOI:** 10.1101/148262

**Authors:** J.A. Habyarimana, E. Baise, C. Douny, M. Weber, F. Boemer, P. De Tullio, T. Franck, C. Marcillaud-Pitel, M. Frederich, A. Mouithys-Mickalad, E. Richard, M.-L. Scippo, D. Votion, P. Gustin

## Abstract

Hypoglycin A (HGA) is a toxin contained in seeds of the sycamore maple tree (*Acer pseudoplatanus*). Ingestion of this amino acid causes equine atypical myopathy (AM) in Europe. Another variety, *A. negundo,* is claimed to be present where AM cases were reported in the US. For unknown reasons, occurrence of this disease has increased. It is important to define environmental key factors that may influence toxicity of samaras from *Acer* species. In addition, the content of HGA in seedlings needs to be determined since AM outbreaks, during autumn period when the seeds fall but also during spring when seeds are germinating. The present study aims to validate a reliable method using high performance thin layer chromatography for determination and comparison of HGA in samaras and seedlings.

The working range of the method was between 20 μg HGA to 408 μg HGA per ml water, corresponding to 12 - 244 mg/kg fresh weight or 40 - 816 mg/kg dry weight, taking into account of an arbitrary average dry matter content of 30%. Instrumental limit of detection and limit of quantification were of 10 μg HGA/ml and 20 μg HGA/ml water, respectively. Instrumental precision was 4% (RSD on 20 repeated measurements) while instrumental accuracy ranged between 86% and 121% of expected value. The HGA recovery of the analytical method estimated from spiked samaras and seedlings samples ranged between 63 and 103%. The method was applied to 9 samples of samaras from *Acer pseudoplatanus, A. platanoides* and *A. campestre* and 5 seedlings samples from *A. pseudoplatanus.* The results confirm detection of HGA in samaras from *A. pseudoplatanus* and the absence of detection in samaras of other tested species. They also suggest that detected levels of HGA are highly variable. This confirmed the suitability of the method for HGA detection in samaras or seedling.

## INTRODUCTION

Hypoglycin A (HGA) is assessed to be a potential inhibitor and a key ingredient involved in the metabolism process of fatty acids. This nonproteinogenic amino acid is contained in ackee (*blighia sapida*), a fruit found in Jamaica [1] and in seeds of some maple species [2], like the sycamore maple tree (*Acer pseudoplatanus*). Ingestion of HGA causes equine atypical myopathy (AM) in Europe [3; 4]. These diseases are characterized by the abrupt symptoms onset resulting from the acute necrosis of postural, respiratory muscles and the myocardium [4]. Horses exhibit depression, weakness, stiffness, recumbency, trembling, sweating and dyspnea [5; 6].

Data from epidemiological studies have shown the seasonality of AM with outbreaks in autumn that are often followed by smaller outbreaks in the subsequent spring [6], resulting from HGA ingestion contained in seeds and seedlings, respectively [7]. All the circumstances leading to AM are not yet fully established, with an increasing number of cases in the recent years [8]. More than 1800 horses throughout Europe have suffered from AM since 2006 (data from the Atypical Myopathy Alert Group website; http://www.myopathieatypique.be), with a mortality rate of 74% [6].

To our knowledge, no curative treatment has been so far documented. Therefore, preventive measures will be of high importance and can only be managed by identifying the sources and the rate of exposure of grazing horses to HGA as well as gaining a good understanding of the factors that influence the concentration of HGA in the horse’s environment. The seasonality of AM outbreak and occurrence of sporadic cases lead us to question the importance of climatic conditions and pasture composition in the development of this disease. These points deserve further investigations to improve preventive measures.

Clarifying the role of maples in AM [9] requires a validated methodology for quantifying HGA in plant extracts. Thus, high performance thin layer chromatography (HPTLC) can be used as a cheap and quick diagnostic tool to measure HGA toxin in a reliable way in maple extracts and useful for valid comparisons between samples collected in a standardized way.

A literature survey concerning HGA problematic revealed that no readily available method, being both practical and cheap, has been developed until now. Indeed, the literature reports conventional chemistry techniques, such as high performance liquid chromatography (HPLC) [10; 11] for determination or quantification of HGA in biological samples [12] in ackee fruit [13], but not in maple samaras, nor in other plant extracts, such as seedlings or leaves. Determination and quantification of HGA in plant extracts using HPTLC, under standardized conditions appeared to be suitable for quick screening. This quantification was done without any special pre-treatment of the sample and using relatively cheap material.

## OBJECTIVES

The present study aims to develop a high performance thin layer chromatography (HPTLC) technique as a cheap and quick diagnostic tool to determine HGA toxin in a reliable way in maple plant material extracts. The technique will be tested on seedlings of *A. pseudoplatanus* and various samaras samples collected on maple trees (*A. pseudoplatanus, A. platanoides* and *A. campestre*).

## STUDY DESIGN

Hypoglycin A was extracted from seedlings and samaras using methanol. After methanol evaporation, the dry extracted was solubilized in water before HPTLC analysis. Different known concentrations of solutions of a commercial HGA standard were used to establish a calibration curve, by plotting the signal intensity against the concentration. Quantification of HGA in plant samples was performed by measuring the signal intensity of the dot on the HPTLC plate corresponding to HGA. This analytical procedure was validated for parameters such as linearity, limit of detection and quantification, precision and accuracy. The analysis of 14 plant samples (supposed to contain or not HGA) was also performed.

## METHODS

### Reagents and preparation of standard solutions

Methanol of analytical grade was obtained from VWR International^®^ (Leuven, Belgium). Water was purified using a Millipore^®^ filtering system (Merck Millipore). S-Hypoglycin A (HGA, 85%) was provided by Toronto Research Chemicals Inc. (TRC^®^, Canada). Ninhydrin (powder, 99%) and acetonitrile of analytical grade (98%) were provided by UCB, Leuven in Belgium.

### Plant Material

Samaras are considered as indehiscent dry fruits called akene. In *Acer*, samaras are associated by pair. Each samara contains one seed. Samaras are especially produced with a pericarp protrusion shaped like a membranous wing. This wing form has a seed spreading function. The shape of the pericarp varies from one species to another. This set (seed and pericarp) is considered as a whole unit and is named “samara” in this study (Fig 1).

**Figure 1:**
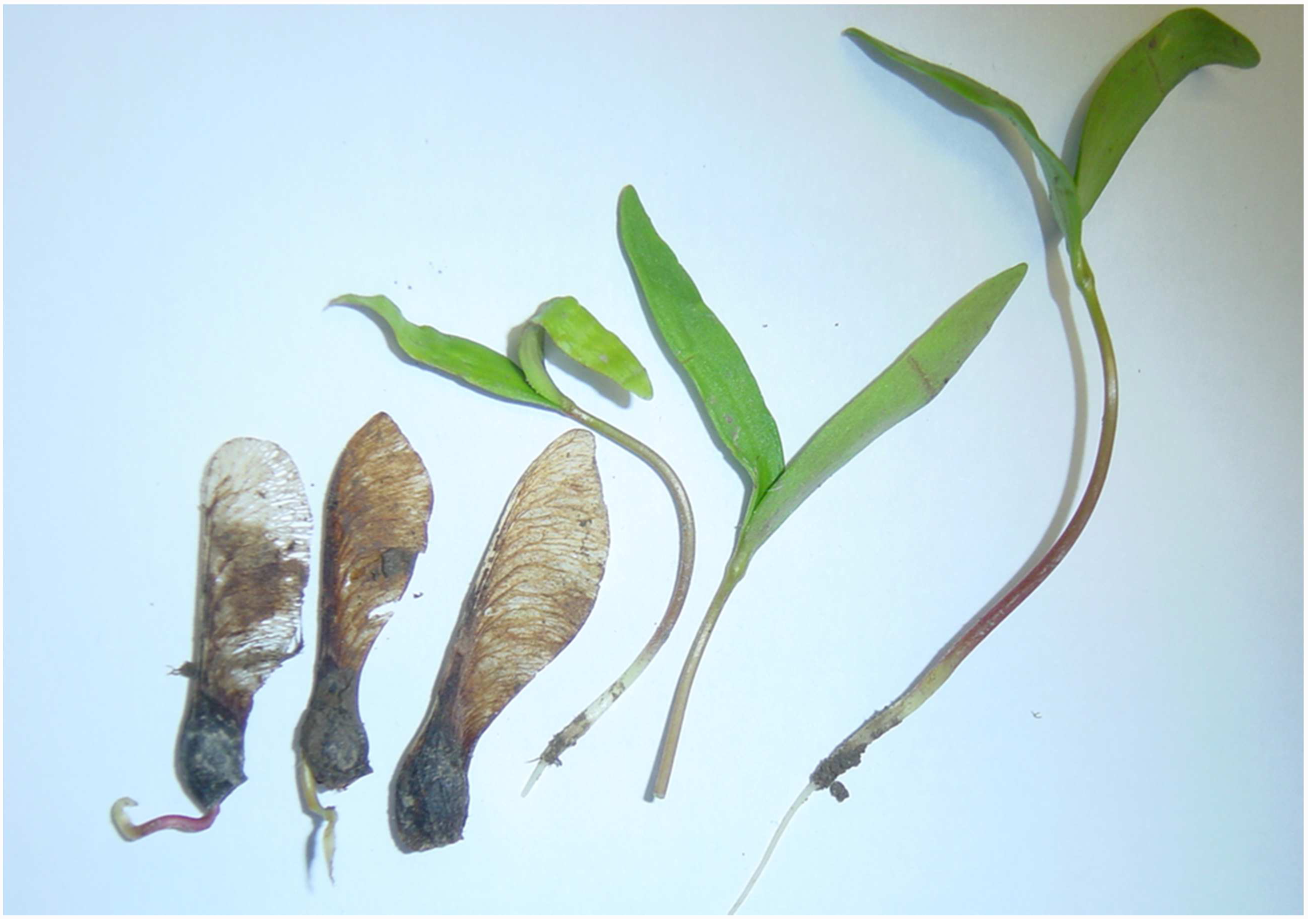
*Acer pseudoplatanus* samara and seedling. Samaras used in this study are complete (third position from left to right) and seedlings are on early two-leaves stage of development (right side of picture).

Specific sites were selected in the areas of pastures where cases of AM were clinically confirmed. Samaras were collected from June to October 2016 directly on the tree and analyzed within 2 days. Seedlings from *A. pseudoplatanus* (2 leaves stage) (Fig 1) were collected from March to May 2016 and stored directly at -80°C until analysis. They were collected around mature identifiable and recognizable trees of the same species. The developed HPTLC method was applied to 9 samples of samaras (five from *A. pseudoplatanus* species, four from *A. plantanoides* and one from *A. campestre)* and five samples of seedlings from *A. pseudoplatanus* at early stage.

### High Performance Thin Layer Chromatography analysis

Chromatographic separation was performed using silica plates (60Å F254, 20 × 10 cm or 10 × 10, Merck Darmstadt, Germany). Three microliters of standards and samples were applied as bands of 4 mm wide, 0.3 mm high and 6 mm apart, using a CAMAG Linomat 5^®^ sample applicator equipped with a 100 μl Hamilton^®^ syringe. Samples were applied at 15 mm from the bottom edge of the chromatographic plate. The plates were allowed to dry for 15 min before elution using a solution of methanol/acetic acid/water (70:20:10, v/v/v). The developing distance was 85 mm, measured from the lower edge of the plates. Post-chromatographic derivatization was obtained by three thin sprays of ninhydrin solution (0.2% in methanol) in 3 different directions [14]. The plate was incubated at 110 °C for 5 min for visualization of the spots. Migration distance was measured and retention factor (RF) was calculated. Chromatograms were obtained by reading the plate at 490 nm using a CAMAG^®^ TLC scanner 3 and the WinCATS^©^ 4.3 software (CAMAG, Muttenz, Switzerland). A negative control (procedure blank) and a positive control (samara sample containing 150 mg HGA/kg fresh weight) are analyzed on each plate. Unknown samples extracts are spotted in triplicate (Fig 2).

**Figure 2:**
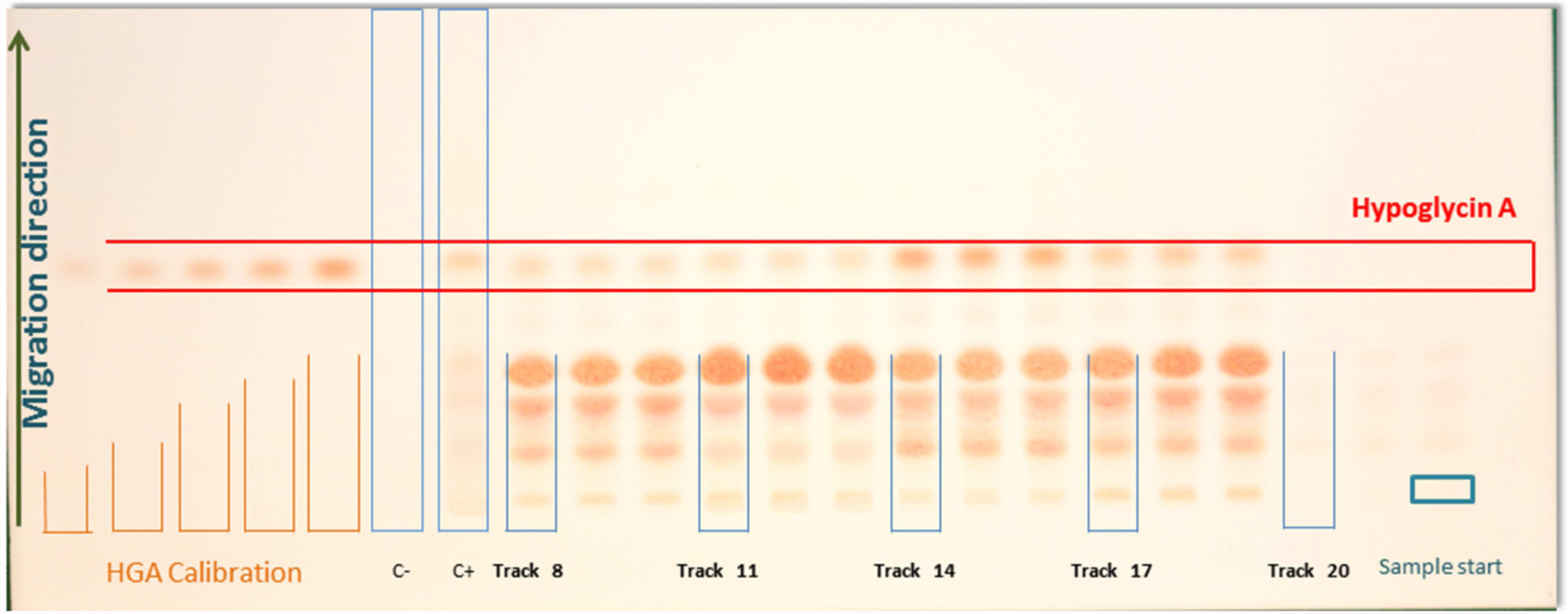
Picture of an HPTLC plate after ninhydrin derivatization and revelation. Tracks 1 to 5: increasing concentrations of hypoglycin A (HGA) in water (20; 40; 85; 199 and 408 μg/ml). Track 6: negative control (C-). Track 7: positive control (samara sample containing 150 mg HGA/kg fresh weight). Track 8 to 22: five samples from samaras of *Acer pseudoplatanus* spotted in triplicate.

### Calibration curve

Ten concentration points were used, fulfilling the criteria required by international conference on harmonization (ICH) concerning the calibration curve [15]. The calibration curve was constructed using HGA solutions in water at respective concentrations of 0, 10, 20, 40, 85, 97, 147, 199, 302, 408 and 510 μg/ml from which 3 μl were loaded on the plate.

### Validation parameters

The analytical procedure was validated according to the guidelines from the ICH of technical requirements for registration of pharmaceuticals for human use [15]. According to these guidelines, the following parameters are to be assessed: specificity, linearity, range, accuracy, precision (intra-day and inter-day precision), detection limit and, quantification limit. In addition, recovery was assessed from HGA spiked samara and seedlings samples.

Specificity was assessed by analyzing procedure blanks (the whole analytical procedure, but without sample). Instrumental precision and accuracy was evaluated by the repeated (n=5) analysis of solutions of HGA in water at 3 different concentrations (57; 177 and 283 μg/ml), on 4 different days. For each level of concentration, precision was expressed as the coefficient of variation (CV) associated to the mean concentration calculated from the repeated measurement, while for accuracy, the mean measured concentration was expressed as a percentage of the theoretical concentration, considered as the “true” value. According to ICH guidelines [16], limit of detection (LOD) was visually evaluated based on calibration curve and selected value tested for reliability. Limit of quantification (LOQ) corresponded to the first point of the calibration curve which was quantified with acceptable accuracy and precision.

Recovery of the method was determined using the standard addition method [17]. Samaras and seedlings (5 g fresh weight), spiked with two amounts of HGA (212 and 408 μg, respectively, corresponding to 127 and 245 mg/kg fresh weight, respectively) as well as unspiked, were extracted as described above (5 repeated analysis). The amount of HGA measured from the unspiked samples was subtracted from the amount measured in the spiked sample. The resulting amount was compared to the spiking level and the recovery was expressed as a percentage of the spiking level.

#### Stability of HGA during the chromatographic analysis

The stability of HGA during the chromatographic analysis was investigated. This was assessed with standard solution of HGA (200 μg/ml) in water and tested by a 2-dimensional HPTLC development. This was performed by two successive HPTLC developments, in 2 perpendicular directions. At the end of these two consecutive operations, components were detected and located in relation to the diagonal axis of the chromatogram (Fig 3). No additional spot was observed.

**Fig 3:**
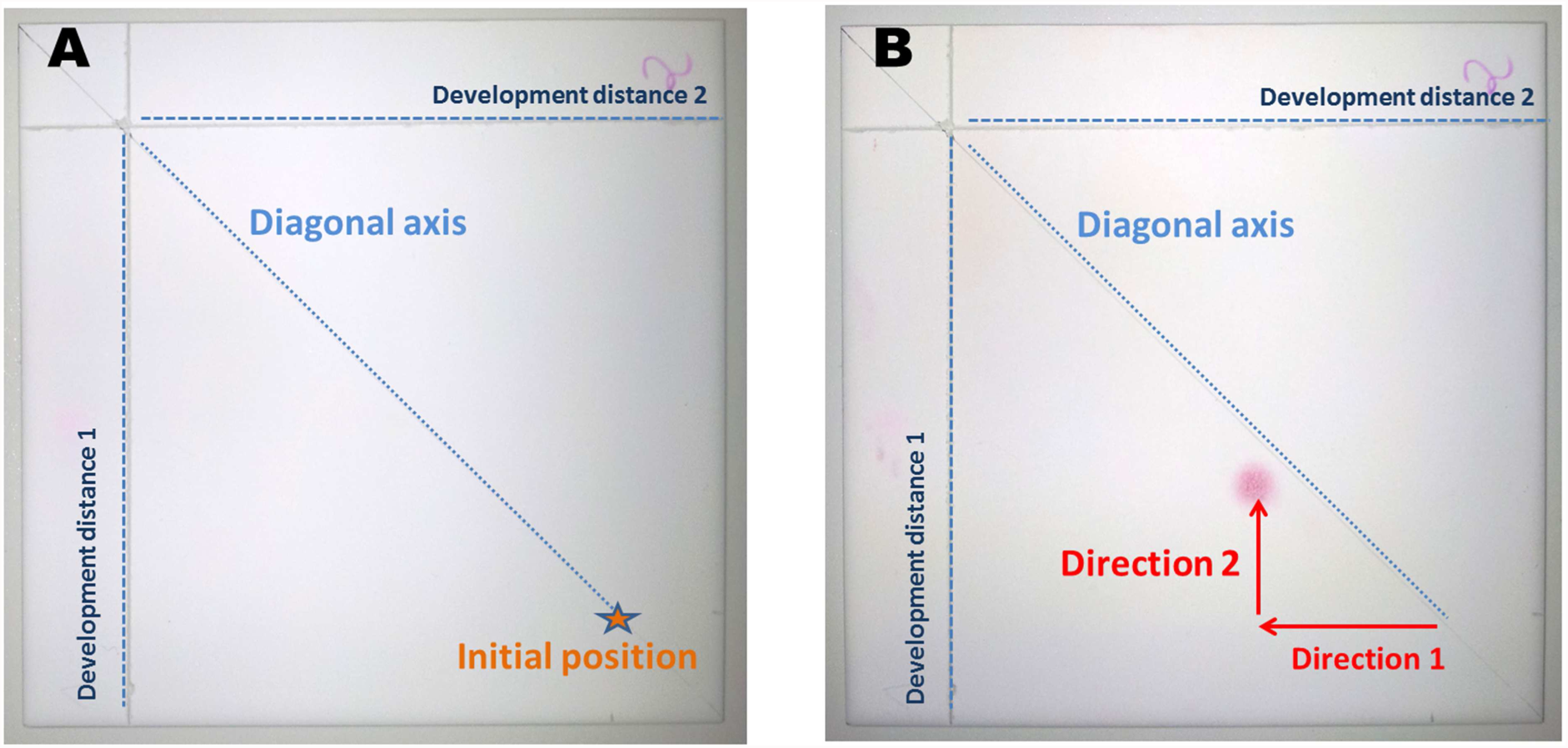
Picture of chromatogram. HP silica plate is used twice successively in two perpendicular directions. The plate is dried before performing the second migration in a perpendicular direction (direction 1 and then 2). Pictures A and B describe the situation before the first development and after the second development, respectively.

### Optimization of extraction procedure

#### Determination of the adequate solvent

Plant material (samara or seedlings) were precisely weighed and crushed. They were mixed with three different solvent (methanol, water or acetonitrile) in three separate sealed vials. A permanent agitation was ensured in order to promote a good contact with solvent. Three arbitrary set extraction periods were planned for solvent collection before analysis of the samples by HPTLC. The concentration of HGA was then calculated and necessary corrections were applied, as for the volume (Fig 4A).

**Fig 4:**
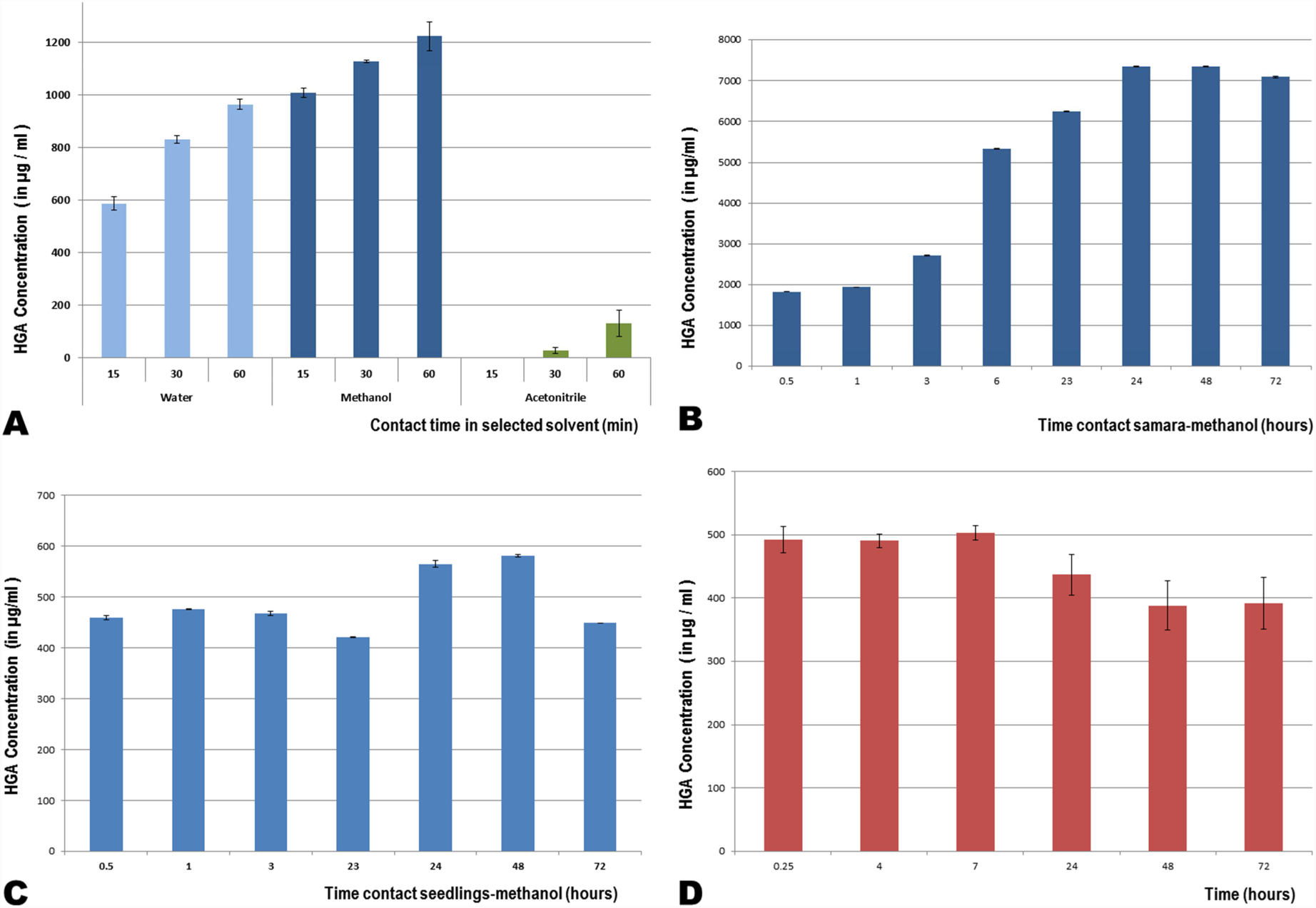
A. Figure summarizing the HGA extraction results on samara, performed on three different solvents (filtered water, methanol and acetonitrile). B. HGA diffusion from samaras. They’re crushed and immersed in methanol. The whole mixture is placed in a tube with constant and moderate agitation. 18 measures of HGA rates are carried over 72 hours. C. HGA diffusion from seedlings. They’re crushed then immersed in methanol. Measures of HGA rates are carried over 72 hours. D. HGA Stability. Standard solution of HGA is placed in a closed tube in laboratory room and under permanent and moderate agitation (same condition as for plant extracts). HGA rate measures are performed during 72 hours.

#### Determination of contact time with adequate solvent

Time contact between plant materials was assessed by placing precise amount of samples in a sealed container in presence of the selected solvent. After determined periods, some solvent was collected and analyzed to determine HGA concentration. Eighteen measures were performed in a period of time spaced from 15 min to 72 hours. Decision was taken to work at laboratory temperature to ensure comfortable and repeatable work conditions and avoid any HGA heat-related alteration (Fig 4 B and C).

#### Stability of HGA during extraction process

A solution of HGA was kept in a sealed vial, in the same conditions than applied with plant material for extraction (laboratory temperature and permanent shaking). Several samples were then collected and analyzed in a period of time from 15 min to 72h (Fig 4 D).

### Determination of the recovery rate

The determination of the HGA recovery rate was performed by using the standard addition method technique [17]. Three increasing amounts of HGA were added to 3 containers, corresponding to three different levels of HGA concentrations, compatible with the HGA concentrations reported in the literature. The analysis was performed in triplicate.

### Extraction of samara and seedlings extracts

Five grams of fresh weight sample were mechanically grinded using a household mixer (Moulinex, Moulinette^®^), were mixed with 25 ml of methanol and extracted during 24 hours at laboratory temperature (about 20°C) under permanent mild shaking. After centrifugation (10000 *g*, 10 min), 12,5 ml of supernatant was evaporated to dryness, and the dry residue was dissolved in 3 ml of water.

For each sample, dry matter was measured independently after 72 hours drying in an oven at 70 °C. The amount of HGA in the sample was expressed in mg/Kg dry matter.

## RESULTS AND DISCUSSION

### Characteristics and validation of the analytical method

HGA showed a RF of 0.51 ± 0.01, determined from the analysis of 10 measurements of HGA solutions (170 μg/ml). Appropriate separation is commonly accepted when 0.1 < Rf < 0.9 [18]. Figure 2 shows a picture of the plate after revelation loaded with a calibration curve (Table 1), a negative (procedure blank) and a positive control and samples from *A. pseudoplatanus.* The procedure blank showed no spot at the RF of HGA, and generated no peak after 490 nm reading, showing the good specificity of the method.

**Table 1:**
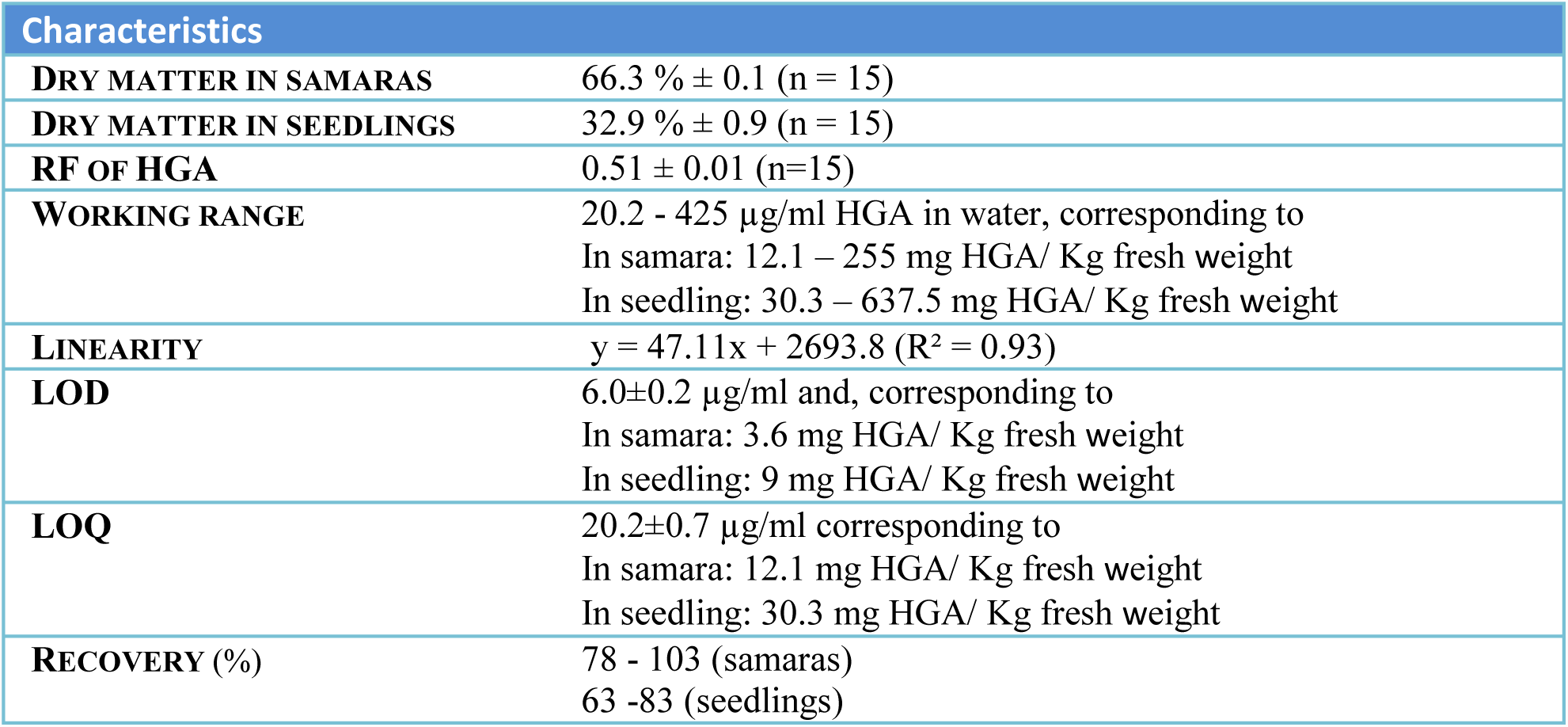
The validation parameters of the HPTLC method for the quantification of HGA in plant extracts. The two values for repeatability and recovery are respectively related to the two mean introduced concentration values.

The validation parameters of the HPTLC analytical procedure are as follow: the dose response relationship was linear in the working range of 20 to 408 μg/ml HGA in water (corresponding 12-245 mg HGA/kg fresh weight or 40-816 mg HGA/Kg dry matter of 30% content). Samples extracts displaying a response above the upper limit of the working range are diluted and reanalyzed.

LOQ was set at 20 μg/ml, corresponding to the fist point of the calibration curve determined with acceptable accuracy and precision (7% and 19% respectively). Accuracy and precsion were of 37% and 16% respectively for the lowest concentration spotted on plate (10μg/ml). This concentration was considered as the LOD. Expressed in mg/kg of plant, the LOD was 6 mg/kg fresh weight and 20 mg/kg dry matter; taken into account of a dry matter arbitrary set to 30% (dry matter content of samara and seedlings is highly variable according to sampling conditions). The LOQ was 12 and 40 mg/kg fresh weight and kg dry matter, respectively. These results are gathered in table 1.

The recovery of the method ranged from 78% to 103% for the levels of concentrations added to samara samples, while 63% to 83% recovery was observed in seedlings. Recovery was tested with HGA concentrations of 42 and 85 mg/kg fresh weight. These levels of recovery were considered as acceptable and consistent with a diagnostic tool to determine the sources of HGA, as targeted in this study.

Intra-day and inter-day instrumental precision and accuracy of the method were calculated (Table 1). Intra-day precision, expressed as a relative standard deviation (RSD), is ranging from 0 to 3%. As expected, inter-day precision is a little bit higher (4%). For accuracy, inter-day accuracy appears to be better (89; 114% and 95% of the introduced HGA concentration for HGA concentrations in water of 57; 177 and 283 μg/ml, respectively) than intra-days accuracies, which ranged from 86 to 121%, depending on the target concentration and the day of analysis. Generally, a range of 80 to 120% of the target value is considered as acceptable for accuracy (Table 2).

**Table 2:**
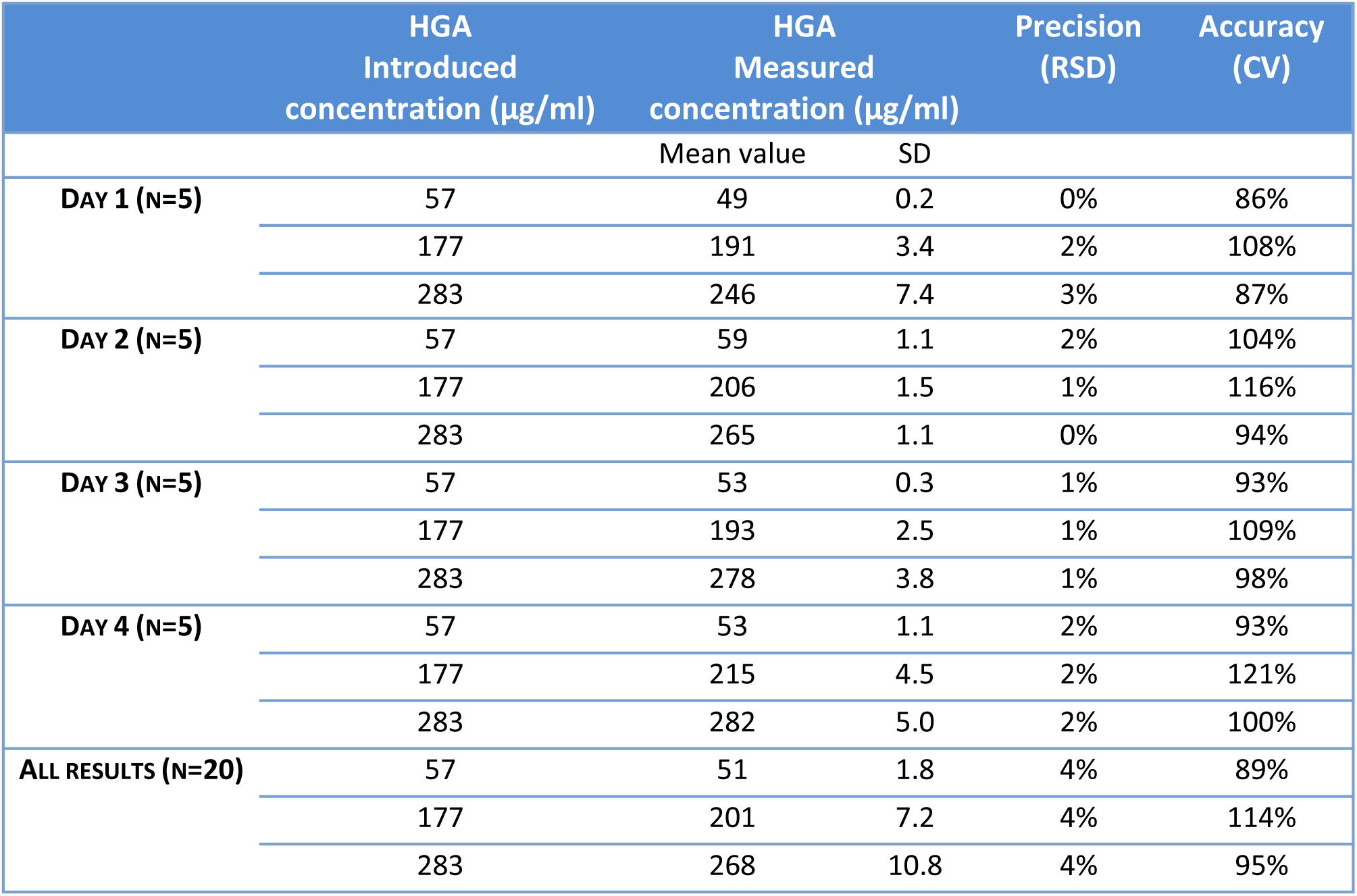
Instrumental precision and accuracy of the HPTLC method to determine hypoglycin-A, calculated from repeated analysis (n=5) of water solution of HGA at 3 concentrations, on 4 days. Precision is expressed as the relative standard deviation (RSD) and accuracy as the ratio between introduced and measured concentration, expressed in percentage (CV).

During routine analysis of unknown samples, a positive control is analyzed on each plate, consisting of a samara sample containing 150 mg HGA/kg fresh weight. The results of the plate are accepted only if the HGA concentration measured in the positive control is in the range 120 - 180 mg/kg fresh weight.

### Analysis of plant samples

Table 3 shows results of HGA concentration determined in samples of samara and seedlings as well as their dry matter content. For Samaras from *A. platanoides* and *A. campestre*, no spot could be observed and no peak could be detected in the area corresponding to the RF expected for HGA. This result is supported by the literature [19]. In Samaras from *A. pseudoplatanus,* HGA concentrations ranged from 678.3 and 918 mg/kg fresh weight (or 2614 to 3104.6 mg/kg dry matter), while in seedlings of the same species, HGA concentrations ranged from 729 and 3028.4 mg/kg fresh weight (or 2563.4 to 11397 mg/kg dry matter). These results show a high variability of HGA concentrations in *A. pseudoplatanus* samaras. Samples were taken at different times through the season, regardless of factors like rain and morning dew. Effects such as seeds leaching or sun drying cannot be seen in our results. The HGA concentrations found in samaras and seedlings in this study are comparable to those found by Unger and co-workers [20], but are slightly lower than those reported by Westermann and collaborators [19].

**Table 3:** hypoglycin A (HGA) concentrations measured in samara and seedling samples (LOD: limit of detection; FW: fresh weight; DM: dry matter).

|  |  | HGA<br>(mg/kg FW) | DRY<br>MATTER<br>(%) | HGA<br>(mg/kg DM) | COLLECTED ON |
| --- | --- | --- | --- | --- | --- |
| **SAMARAS**<br>*A. pseudoplatanus* | *Pseudoplatanus 1* | 678.3 | 23% | 2950.6 | July 2016 |
|  | *Pseudoplatanus 2* | 785.5 | 25% | 3104.6 | July 2016 |
|  | *Pseudoplatanus 3* | 918.0 | 30% | 3095.0 | August 2016 |
|  | *Pseudoplatanus 4* | 797.0 | 30% | 2614.0 | August 2016 |
| **SEEDLINGS**<br>*A. pseudoplatanus* | Seedlings 1 | 2162.3 | 25% | 8798.2 | March 2016 |
|  | Seedlings 2 | 3028.4 | 27% | 11397.0 | March 2016 |
|  | Seedlings 3 | 911.8 | 13% | 6876.1 | April 2016 |
|  | Seedlings 4 | 729.0 | 28% | 2563.4 | May 2016 |
|  | Seedlings 5 | 767.6 | 21% | 3664.1 | May 2016 |
| **OTHER SAMARAS** | *Platanoides 1* | < LOD | 49% | < LOD | Oct. 2016 |
|  | *Platanoides 2* | < LOD | 46% | < LOD | Oct. 2016 |
|  | *Platanoides 3* | < LOD | 54% | < LOD | Oct. 2016 |
|  | *Platanoides 4* | < LOD | 53% | < LOD | Oct. 2016 |
|  | *Campestre* | < LOD | 44% | < LOD | Oct. 2016 |

## MAIN LIMITATIONS

No harmonized measure allows easy comparison of different studies, over a wide variety in results, whether between different plant species or within the same group, or even because of collecting season. Laboratory practice also matters: considering the whole samara or selected fragments allow to express results in so little comparable way. The main limitation of the method lies in the fact that despite its precision it does not give the amount of HGA ingested by the animal but provides an indicative value of its availability in the environment of the horse.

## CONCLUSIONS

A simple, cost effective, specific and accurate HPTLC method for detection, determination and quantification of HGA in maple extracts from *A. species* has been developed and validated. This standardized method was shown to be applicable as well on samaras as on seedlings.

It is also conceivable that with some proper adjustments and calibration, our method can quickly be adapted for other vegetal samples as well as leaves or stems or adapted to other vegetal species as grass for example. This precise and flexible method, easily adaptable, brings a valuable input and can become an interesting tool in the research for a cheap, rapid test apparatus in the diagnosis of AM or seasonal pasture myopathy resulting from HGA intoxication in horse.

## ACKNOWLEDGMENTS

Dr Ursula Fogarty from the Irish Equine Center is gratefully acknowledged for advices regarding the manuscript. The study was supported by the “Institut français du cheval et de l’équitation (*If*ce)” of France and by “Les Fonds Spéciaux pour la Recherche (FSR)” of Liege University (Belgium).

Conflict of Interest: authors disclose no conflict of interest.

